# Proliferation specific codon usage facilitates oncogene translation

**DOI:** 10.1101/695957

**Authors:** Hannah Benisty, Marc Weber, Xavier Hernandez-Alias, Martin H. Schaefer, Luis Serrano

## Abstract

Tumors evolve under selection for gene mutations that give a growth advantage to the cancer cell. Intriguingly, some cancer genes are more often found mutated in tumors than their closely related family members. For example, KRAS mutations are more frequently observed in cancer in comparison to HRAS and NRAS. Here, we find that for RAS and six oncogene families, the most prevalent mutated members in cancer have a codon usage characteristic of genes involved in proliferation. The codon usage of KRAS is more adapted to be efficiently translated in proliferative cells than the codon usage of HRAS. We also show that the translation efficiency of KRAS varies between cell lines in a manner related to their tRNA expression. Altogether, our study demonstrates that a dynamic translation program contributes to shaping the expression profiles of oncogenes. We propose that codon bias related to cell proliferation contributes to the prevalence of mutations in certain members of oncogene families.

## Introduction

Cancers arise due to mutations that confer a selective growth advantage on cells (Nowell, 1976). These mutations can occur in oncogenes, which when activated by mutations contribute to the cancer proliferation phenotype. Interestingly, oncogenes often have closely-related family members that are less frequently mutated in cancer.

The RAS family is a striking example. Activating mutations in KRAS are among the most common mutations in human cancers (Lawrence et al., 2014). KRAS belongs to a family of three genes, the other two being HRAS and NRAS. The encoded proteins share a high sequence identity of 85% and hence similar structure and biochemical properties (Hobbs et al., 2016). However, the reasons for the drastic variation in mutation incidence between the RAS genes remain enigmatic. Significant effort is being invested in studying the molecular differences between the three family members and specifically to understand what is special about KRAS (Drosten et al., 2017); (Haigis et al., 2008); (Quinlan and Settleman, 2008); (Lampson et al., 2013); (Koera et al., 1997; Potenza et al., 2005); (Omerovic et al., 2008; Yan et al., 1998); (Apolloni et al., 2000; Rocks et al., 2006); (Prior et al., 2003)). An intriguing observation is that even though RAS proteins are very similar, the codon usage is different, with only 15% of codon identity (Lampson et al., 2013). The nucleotide sequence of KRAS is enriched in rare codons (decoded by low-abundant tRNAs) in comparison to HRAS. This has been linked to a poor translation efficiency of KRAS and a high efficiency for HRAS (Lampson et al., 2013). It has been speculated that these differences in codon usage relate to the imbalance of the mutation frequency within the RAS family: the constitutively activated form of the highly translated HRAS might lead to an over-activation of the MAPK pathway, ultimately leading to oncogene-induced senescence (Bodemann and White, 2013; Pershing et al., 2015).

Codon usage and tRNA abundance are important parameters for fine-tuning protein synthesis. The functional influence of codon optimality and tRNA levels on the efficiency of protein production remains a topic of intense debate (Hanson and Coller, 2018). In recent years, studies have shown that tRNA levels are not static but dynamically regulated in different cellular contexts, leading to changes of the translation efficiency of transcripts depending on their codon composition (Torrent et al., 2018; Gingold et al., 2014; Goodarzi et al., 2016). In mammalian cells, changes in tRNA abundance have been reported across different cell states, and specifically between healthy and cancer cells (Gingold et al., 2014; Goodarzi et al., 2016). Interestingly, Gingold et al (Gingold et al., 2014) showed that a specific subset of tRNAs are up-regulated in proliferating cells, while they are downregulated in differentiated or arrested cells. Additionally, they show that genes that are necessary for cell division have a codon usage adapted to the tRNA repertoire in proliferative cells. Thus, changes in the expression of specific tRNAs could regulate an entire functional class of genes, for instance proliferative genes, to favor cell growth. Would a cancer cell take advantage of this translational program to modulate the expression of cancer genes to its own growth advantage? Could it be that a dynamic regulation of RAS translation efficiency determines the uneven mutation frequencies across RAS genes? Will this be a general phenomenon in other cancer gene families?

To answer the above questions we first identify 8 protein families of three members, RAS, RAF, RAC, RHO, FOXA, FGFR, COL and AKT, with high protein sequence similarity and at least one protein being relevant for cancer. We find that in all but one family the codon usage signature of the most frequently mutated gene is characteristic of proliferation-related genes in comparison to its homologous family members. We then study the RAS family composed of KRAS, HRAS and NRAS in detail. We measure how proliferation and quiescent cell states induce codon-dependent changes in KRAS protein levels. Finally, we find that different tRNA expression profiles between cell lines correspond to differences in KRAS protein levels. This work supports the existence of different translational programs such as the up-regulation of proliferative tRNAs that have the potential to boost the protein synthesis of oncogenes. Thus, our results suggest that dynamic changes in this fundamental cellular process may contribute to cancer and specifically to the prevalence of mutations in certain genes as compared to their closely-related family members.

## Results

### Codon usage of cancer genes

To explore whether the differences in mutation frequency between RAS genes are also observed in other gene families, we perform a genome-wide survey in a pan-cancer data set from The Cancer Genome Atlas (TCGA) to identify gene families with variation in mutation incidence in cancer. To define families, we cluster sets of proteins based on protein sequence similarity. We restrict the analysis to families containing at least one known cancer driver gene (Lawrence et al., 2014). We identify 8 families including the RAS family. We consistently observe one gene more frequently mutated (non-synonymous mutation counts) in comparison to the other genes of the family (Figure 1A, Table S1). Especially, for the RAS, RAF and RAC families we observe at least a two-fold variation in the mutation count number (fold change between the family member with the lowest mutation count and the highest). For the RHO, FOXA, FGFR and COL families we observe between 1.30 to 1.95-fold change. However, this effect is mild for the AKT family with only a 1.22-fold change.

**Figure 1:**
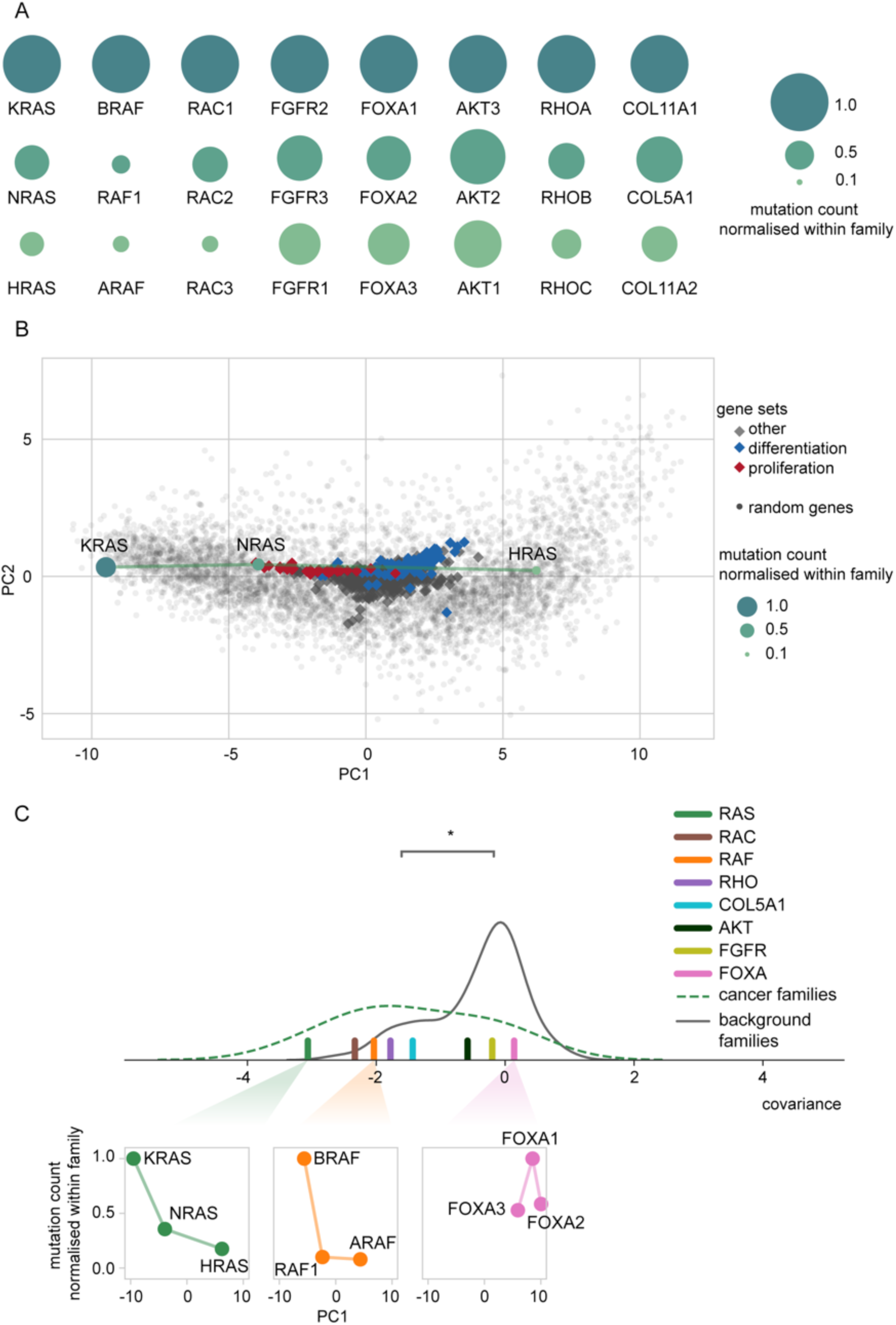
Association between codon usage and mutation frequency in genes from 8 different families. See also Figure S1 and Tables S1. A. Gene triplets with divergent mutation frequencies in cancer. B. PCA projection of the human codon usage. The location of each gene is determined by its codon usage. Distribution of GO gene sets along the main codon usage axis reveals the two functional poles, “proliferation” (negative PC1) and “differentiation” (positive PC1). The positions of the RAS genes and their normalized mutation count are shown. See also Figure S1B. C. Distribution of the covariance of mutation count normalized within family and PC1 (lines are kernel density estimates as a guide to the eye). The covariance of cancer gene families is significantly more negative than the background families (W.M.W. test p<0.018). All families but one (FOXA) have a negative covariance. See also Figure S1C.

As previously described, RAS genes have a high amino acid sequence identity (85%) but differ in their codon usage (15% codon identity) (Lampson et al., 2013). The same observation applies to the 7 other families we selected (Figure S1). This raises the question of whether differences in the mutation count could be related to variation in codon usage in addition to potential biochemical differences on the protein level.

Therefore, we investigate whether the codon usage of these genes could be related to a specific translation program. Previously, Gingold et al (Gingold et al., 2014) described the average codon usage bias in different gene functional groups and observed that genes in two cellular programs, differentiation and proliferation, preferentially use different synonymous codons. Additionally, they found that tRNAs induced during proliferation correspond to the codons that are enriched in the functional set of proliferation genes.

To test if functional adaptation to these cellular programs could have shaped the codon usage in the selected gene families, we examine how the codon usage of the genes correlates to the codon usage of proliferation-related and differentiation-related genes. We use a similar approach to Gingold et al (Gingold et al., 2014) by applying PCA to the relative codon usage frequencies of all individual genes, in order to visualize how the codon usage of the genes of the 8 families correlate with the codon usage of pro-proliferative genes. By computing the projection of all major gene sets in the Gene Ontology (GO) classification, we reproduce the results of Gingold et al (Gingold et al., 2014) revealing two distinct functional poles at the extremes of the codon usage main projected axis, the first principal component (PC1) (Figure 1B). At one extreme, for negative values of PC1, we find a strong enrichment of gene sets that are descendants of the “cell cycle” term (16 out of the top 30, Fisher exact test two-sided p < 2.2e-17). At the other extreme, for positive values of PC1, a majority of the gene sets are descendants of the “multicellular organism development” or “cell differentiation” terms (14 out of the top 30, Fisher exact test two-sided p < 5.8e-6). This observation, together with the previously described changes in tRNAs in proliferative versus non-proliferative cells (Gingold et al., 2014), shows that the two poles of codon usage correspond to two cellular translation programs. We next calculate the average codon usage of each coding sequence of the selected cancer gene families and project it in the PCA plane (Figure S1) as well. We observe that the transcript of KRAS is composed of codons more frequently used by genes involved in proliferation in comparison to HRAS (Figure 1B). This seems to be a general phenomenon as the codon usage of the most frequently mutated family member corresponds better to the codon usage of pro-proliferative genes than their cognate family members, except for the FGFR and FOXA families. FGFR1 and FGFR2 have an inverse relationship, with the gene less often mutated being the one having a pro-proliferation codon usage. In cancer, amplifications are the most common alterations of FGFR1 whereas FGFR2 harbors activating mutations (Sobhani et al., 2018). Thus, in our analysis it is difficult to discern which of the two is more oncogenic. FOXA genes show a codon usage in the opposite pole of the pro-proliferation codon usage (Figure S1B). In this family the cancer driver gene FOXA1 can take the role of a tumor suppressor (Barbieri et al., 2012); (Schroeder et al., 2014), typically inactivated by mutations in contrast to the other families where the most frequently mutated gene is an oncogene with activating mutations. This suggests that the usage of proliferation-associated codons in cancer genes is a characteristic property of oncogenes.

Next, we seek to assess the significance of the correlation pattern between codon usage and mutation frequency. Our main observation is that the gene member that is the most frequently mutated is the one that presents a codon usage most adapted to the proliferation codon usage pole (negative pole of PC1). Thus, we expect that PC1 and mutation frequency are negatively correlated. For the 63 gene families that do not contain any cancer driver gene (non-cancer gene families), we assume that there is no specific relationship between codon usage and mutation frequency, such that the correlation should be randomly distributed around zero (Figure S1C). We also assume that the pattern is more significant when, within a cancer gene family, we observe both a large variation in codon usage and in mutation frequency. Thus, we compare the distribution of the covariance of PC1 and mutation frequency for cancer gene families to the background gene families (Figure 1C, Table S2). The covariance tends to be large, and thus gives more weight to the families that present a large variation in codon usage and in mutation frequency. Families with little variation in either codon usage or mutation frequency, on the other hand, present a smaller covariance. We observe that the covariance of cancer gene families is significantly more negative than the background families (Wilcoxon-Mann-Whitney (W.M.W.) test p<0.018). In particular, 7 out of 8 families (RAS, RAF, COL, RAC, RHO, AKT and FGFR) present a pattern of negative correlation, with the families showing the highest covariance being RAS, RAF, RAC, RHO and COL.

### Codon usage-specific changes of KRAS protein abundance under different cell states

The above analysis suggests that oncogenes with a codon usage signature characteristic of proliferation-related genes will be more expressed under a proliferative cell state. To test this hypothesis, we decide to work with the RAS family.

In order to determine if KRAS protein abundance changes in different cell states in a codon-dependent manner, we establish a series of manipulated cells that co-express KRAS wild-type (KRAS_WT_) and a protein sequence-identical KRAS variant but with a codon usage similar to HRAS (Lampson et al., 2013) (KRAS_HRAS_). The two proteins have different N-terminal tags that allow us to distinguish between the two versions of KRAS by their size (FLAG and 3xHA; as a control we also made an identical construct with the tags swapped Figure S2A). A bidirectional symmetrical promoter controls the simultaneous expression of the two genes. This design provides us with a controlled expression system to assess exclusively codon-dependent changes in protein abundance while reducing the impact of other factors (eg. transcriptional efficiency or biochemical properties of the protein). Moreover, both genes are in the same plasmid and are therefore integrated into the genome with equal stoichiometry (Figure 2A).

**Figure 2:**
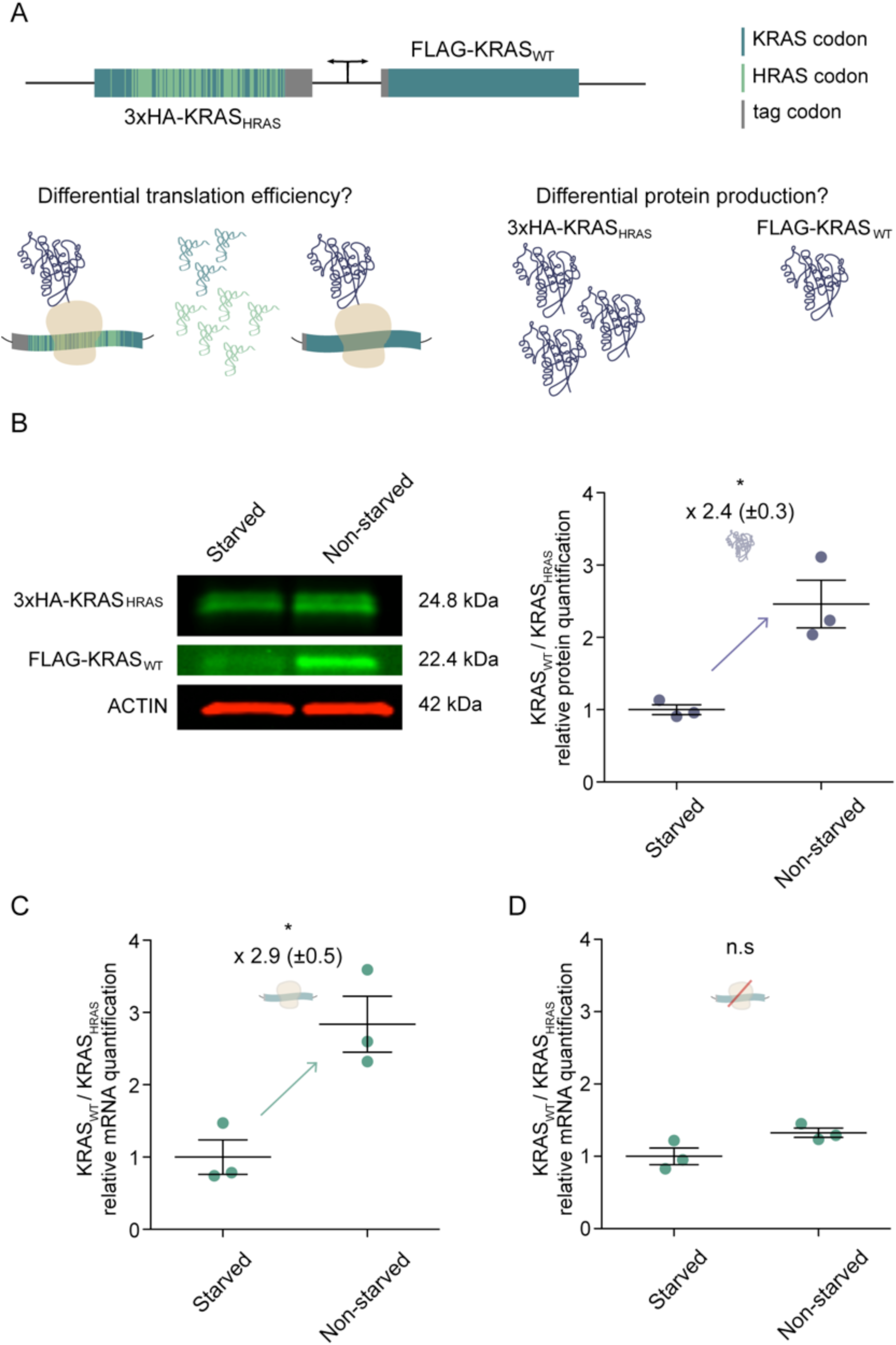
The codon usage of KRAS_WT_ is adapted to be efficiently translated during proliferation. A. Experimental design. The construct co-expresses two genes coding for the same KRAS protein, differentiated by size with two different tags. HRAS-specific codons are represented in green. KRAS-specific codons and identical codons between KRAS and HRAS are represented in blue. KRAS protein is represented in dark purple. Distinct tRNAs pools appear in green and blue. B. Western blot analysis of the levels of KRAS_WT_ and KRAS_HRAS_ in starved and non-starved BJ/hTERT cells. The protein ratio KRAS_WT_/KRAS_HRAS_ increases from quiescent to proliferative state. C. The transcript ratio KRAS_WT_/KRAS_HRAS_ increases between the two cell states. D. Translation inhibition with RBS and ATG site removal decreases the effect on transcript level. Results in B, C and D are representative of three independent experiments with three technical replicates each. Values are relative to starved condition. Error bars represent SEM. * p < 0.05 (unpaired Student t test).

Gingold et al (Gingold et al., 2014) reported changes in tRNA profiles of BJ/hTERT fibroblast in different cell-states: a quiescent state when the cells are starved and a proliferative state when the cells are not starved. Therefore, we first co-express KRAS_WT_ and KRAS_HRAS_ in BJ/hTERT fibroblasts and quantify the protein ratio of KRAS_WT_/KRAS_HRAS_ in these two different cell-states. We observe that the ratio increases by more than two fold when the cells are proliferating (Figure 2B). The observed fold change suggests that KRAS codons are more efficiently translated during proliferation than HRAS codons.

We also measure the ratio at the transcript level and, interestingly, we find the same effect as observed at the protein level: the ratio between KRAS_WT_ and KRAS_HRAS_ is increased by more than two fold in proliferation versus starvation (Figure 2C). Previous studies have shown in different species that codon optimality has a high impact on transcript stability (Presnyak et al., 2015; Boël et al., 2016; Wu et al., 2019). An interesting hypothesis is that the dynamics of ribosomal elongation influences mRNA decay. Ribosome translocation is slower through non-optimal transcripts and promotes mRNA decay, mediated by the DEAD-box protein Dhh1p in *S. cerevisiae* (Radhakrishnan et al., 2016). Thus, codon content directly modulates both translation efficiency and mRNA stability. Our study suggests that KRAS_WT_ is composed of codons that are optimal for its expression in proliferative cells but that are non-optimal in starved cells. Therefore, to determine if changes in KRAS transcript abundance are due to differences in translation efficiency and not transcriptional regulation, we prevent translation by deleting the ribosome binding site and the ATG start codon. We first test to confirm that there is no protein expression when cells are established with the non-productive expression cassette (Figure S2B). After blocking the translation of the two genes we observe that the difference of KRAS_WT_/KRAS_HRAS_ between non-starved and starved is not significant at the transcript level (Figure 2D). In short, KRAS_WT_/KRAS_HRAS_ changes are mainly due to a differential translation efficiency (that also increases the corresponding mRNA level) between a quiescent state and a proliferative state. Our results provide new evidence supporting the dynamic translational efficiency by cell-state-specific codon usage of transcripts.

### Specific differences in tRNA levels explain differences in KRAS abundance between cell lines

To investigate if the condition-specific translation efficiency is mediated by differential tRNA expression, we explore the effect of cell line-specific tRNA abundances on KRAS expression. A previous study (Fu et al., 2018) has already reported a cell line-specific expression of KRAS_WT_ and KRAS_HRAS_. We therefore hypothesise that the tRNA content of different cell lines varies and may influence the translation efficiency in a codon-dependent manner. To test our hypothesis, we first establish two additional cell lines (HEK293 and HeLa) to co-express KRAS_WT_ and KRAS_HRAS_. We verify that the expression results in changes of both protein and mRNA when comparing BJ/hTERT, HEK293 and HeLa (Figure 3A, 3B). Of these three cell lines HEK293 exhibits the highest proliferative rate (Figure S2C) and higher abundance of KRAS_WT_ in comparison to HeLa and BJ/hTERT. We observe the same effect on protein level when switching the position of the tags (Figure S3A), showing that FLAG and 3xHA are not influencing our observation. As before, the removal of the ribosome binding site and start codon leads to similar transcript levels for the three cell lines (Figure 3C), indicating that translation is an important determinant of mRNA stability.

**Figure 3:**
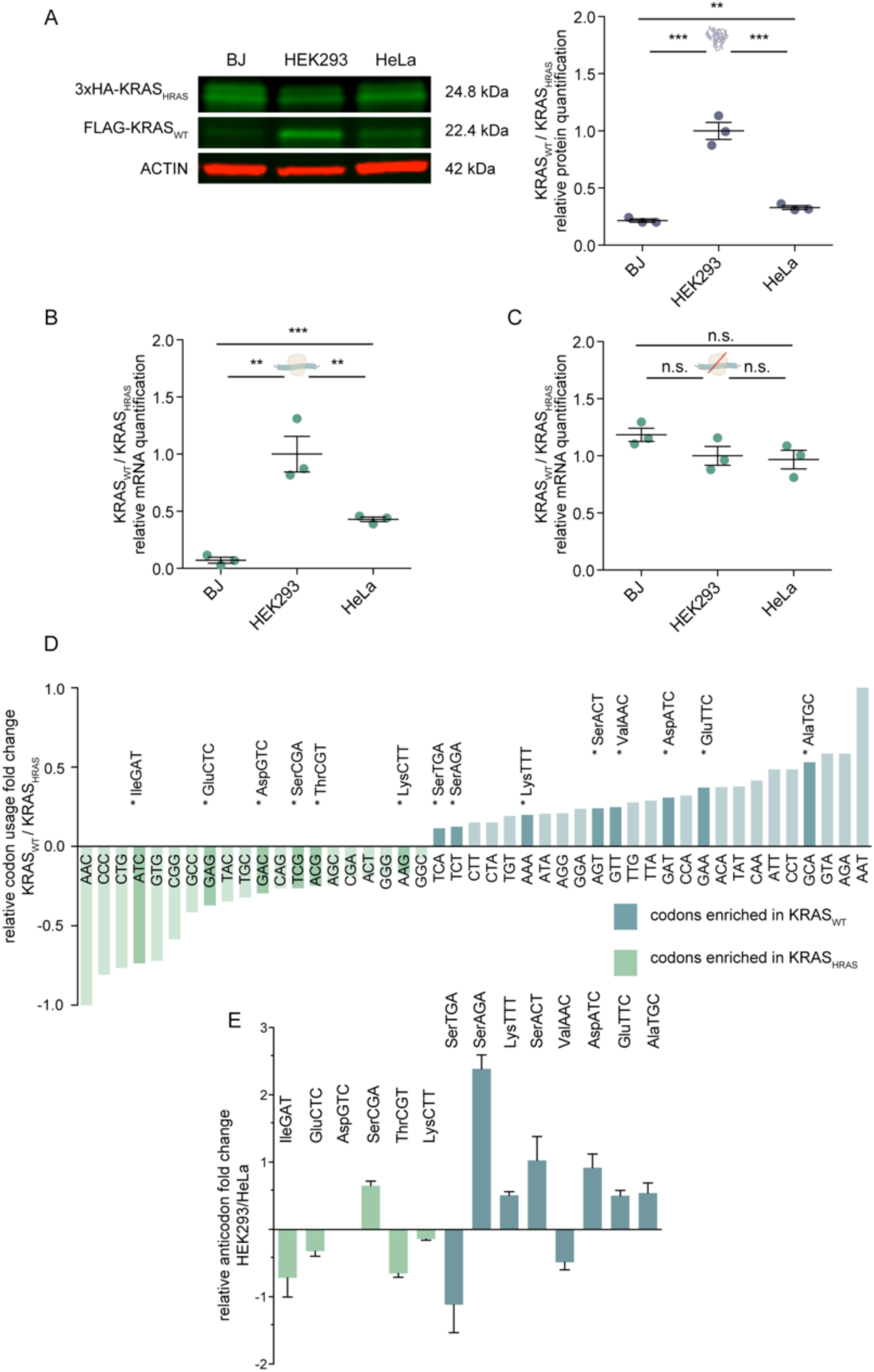
tRNA levels variations are associated to KRAS_WT_ and KRAS_HRAS_ abundance in HEK293 and HeLa. See also Figure S2 and S3 and Tables S3 and S4. A. Western blot analysis of the levels of KRAS_WT_ and KRAS_HRAS_ in BJ/hTERT, HEK293 and HeLa cells. The protein ratio KRAS_WT_/KRAS_HRAS_ varies between the different cell lines. B. The transcript ratio KRAS_WT_/KRAS_HRAS_ also varies between the cell lines. C. Translation inhibition decreases the differential effect observed in cell lines on transcript level. Results in B, C and D are representative of three independent experiments with three technical replicates each. Values are relative to HEK293. Error bars represent SEM. * p < 0.05; ** p < 0.01; *** p < 0.001 (unpaired t test). D. Fold change of the relative codon usage (pseudocount +1) between KRAS_WT_/KRAS_HRAS_. The codons that are not changing in amount between KRAS_WT_ and KRAS_HRAS_ are not represented. tRNAs differentially expressed between HEK293 and HeLa are highlighted. E. Fold change of tRNA expression between HEK293 and HeLa is represented for the cognate tRNAs of the codons enriched in either KRAS_WT_ (blue) or KRAS_HRAS_ (green). Error bars represent SEM of three independent hydro-tRNAseq experiments. Differences between HEK293 and HeLa were assessed by a multiple t test using a permutation based FDR cut-off of p<0.05. * p < 0.05.

The above observations suggest that the effect of codon bias may be differentially regulated in different cell types. If translational efficiency is different in each cell type, we hypothesise that it should match the cell type’s tRNA anticodon abundance. More specifically, we expect that the relative synonymous codon frequencies (relative to the amino acid) of KRAS_WT_ match better to the relative abundances of cognate tRNAs in HEK293 than in HeLa. To associate the amount of tRNAs with codon usage, we perform hydro-tRNA sequencing (Gogakos et al., 2017) and quantify tRNA expression in HEK293 and HeLa cells (Table S4).

We find 14 tRNAs showing significant differences (q < 0.05, t test) between the two cell lines (Figure S2D, Table S4). Six of them are expressed higher in HEK293 and match codons enriched in the coding sequence of KRAS_WT_ (TCT, AAA, AGT, GAT, GAA, GCA). One exception occurs with tRNASer^CGA^, which is expressed higher in HEK293, but the associated codon TCG is not enriched in KRAS_WT_. On the other hand, 5 tRNAs expressed significantly higher in HeLa correspond to codons enriched in HRAS and therefore in KRAS_HRAS_ (ATC, GAG, GAC, ACG and AAG). Only two tRNAs higher in HeLa (tRNASer^TGA^ and tRNAVal^AAC^) do not match codons enriched in KRAS_HRAS_ (Figure 3D-E). To sum up, we find 11 out of 14 tRNAs matching the expected codons in KRAS_WT_ and KRAS_HRAS_. Therefore, the difference in tRNA supply between cell lines could explain the observed variation of KRAS_WT_ and KRAS_HRAS_ protein levels. In a previous study in which different codons of KRAS had been changed, it was observed that certain replacements resulted in significant increases in KRAS expression reaching in a cumulative manner the levels of HRAS (Lampson et al., 2013). Among them were the changes GCA to GCC, AAA to AAG and ATT to ATC, which correspond to the anticodons of tRNAs differentially expressed between HEK293 and HeLa. The changes GAA to GAG and CCT to CCC did not display protein abundance changes (Figure S2E).

Additionally, we investigate if the codons corresponding to the significantly changing tRNAs are also found enriched in the most prevalent oncogenes of the RAF, RAC, RHO and COL families. Overall, the codons enriched in KRAS_WT_ and having their matching tRNAs significantly increased in HEK293 are also enriched in the oncogenes BRAF, RAC1, RHOA and COL11A1 in comparison to their less mutated family member. Conversely, the codons enriched in KRAS_HRAS_ and the matching tRNAs in HeLa, are also enriched in the less frequently mutated genes, RAF1, RAC3, RHOC and COL11A2 (Figure S3). Finally, we confirm that 6 out of 7 tRNAs more highly expressed in the most proliferative cell line HEK293, correspond to proliferation-related codons (Figure 4). Altogether, our results support a dynamic translational program, where specific changes in tRNA abundance can shape the expression of proliferation-related transcripts.

**Figure 4:**
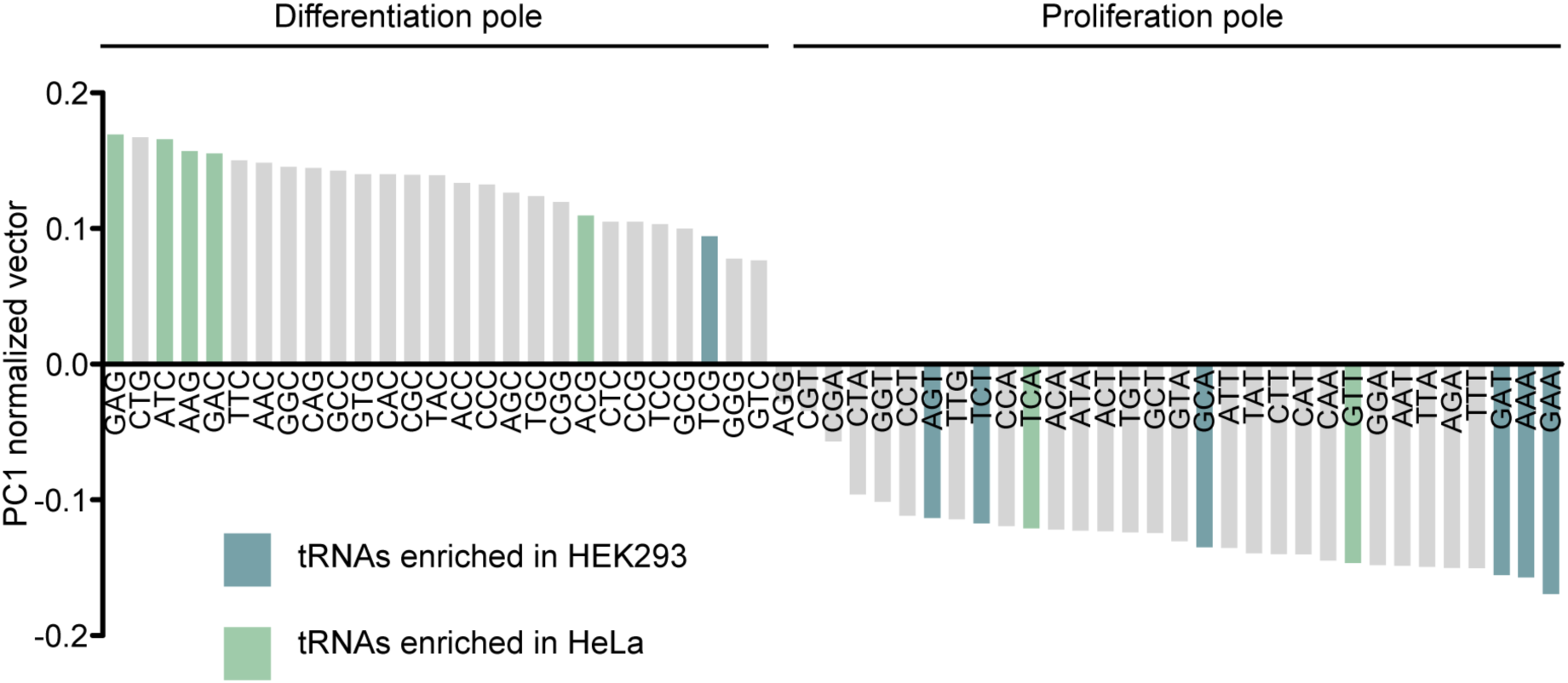
Proliferation-versus differentiation-related codons. See also Figure S3. Codons are ordered following their value in the first component axis (PC1 axis). Note that the scale of the values is arbitrary, only the relative values are important (direction of the vector in the multidimensional space). Negative values indicate negative PC1 toward the proliferation pole, positive values toward the differentiation pole.

## Discussion

Codon usage and tRNA abundance are crucial for efficient and accurate translation of mRNAs into proteins. Previous studies have found that tRNA repertoires are dynamic in a manner that facilitates selective translation of specific transcripts (Gingold et al., 2014; Supek, 2016; Newman et al., 2016; Torrent et al., 2018. Here, we investigate if there is an oncogenic translation program shaping the abundance of cancer driver genes. We describe protein families with strong differences in codon usage and mutation frequencies within the family. The observed codon bias reveals a proliferation-specific codon usage of the more prevalent family members in cancer. Specifically, the families RAS, RAF, RAC, RHO and COL exhibit the largest negative covariance between mutation frequency and proliferation-associated codon usage. This raises the question of whether these transcripts are more effectively translated in proliferative cells than their closely related family members.

We focus on the example of the RAS family and we experimentally show that the translation efficiency of KRAS_WT_ is up-regulated in proliferative cells in comparison to the translation efficiency of KRAS_HRAS_. Additionally, we find that translation efficiency is a determinant of transcript abundance. This observation has been previously described in *H*.*sapiens* (Wu et al., 2019), *S. cerevisiae* (Presnyak et al., 2015) and *E. coli* (Boël et al., 2016). Here, we consistently show that the changes in KRAS_WT_/KRAS_HRAS_ transcript abundance between cell types and cell states decrease when translation is suppressed.

Activating mutations of oncogenes are a product of selection during tumor initiation for an ideal level of signaling. It is plausible that selection acting on a gene depends on the level of expression of that gene. Pershing et al observed that replacing KRAS codons with HRAS codons in one exon renders mice more resistant to lung tumors and decreases the amount of KRAS mutations (Pershing et al., 2015). This supports our hypothesis that translation efficiency might contribute to mutation frequency differences between genes.

RAS abundance is an important determinant of MAPK signaling which is tightly connected to cancer growth. Importantly, it has been shown that cells initially expressing low levels of oncogenic RAS only progressed into malignant lesions after RAS levels increased (Sarkisian et al., 2007; Ferbeyre, 2007). In line with this model, it is tempting to speculate that mutated RAS increases its own translation by triggering cell proliferation (Figure S4).

Our observations are in agreement with recently identified alterations in transcript-specific translation that emerge as drivers of cellular transformation. For example, it has been shown that up-regulation of specific tRNAs (tRNAGlu^TTC^ and tRNAArg^CCG^) in metastatic cells leads to an increase in the amount of certain proteins, specifically EXOSC2 and GRIPAP1 that play an important role in metastasis (Goodarzi et al., 2016). Indeed, we have seen similar results here for KRAS. Consistent with previous reports (Dittmar et al., 2006; Pavon-Eternod et al., 2009), we observe that specific tRNAs vary between different cell lines, which could explain the differences in KRAS expression between HEK293 and HeLa. One of them, tRNAGlu^TTC^, is also upregulated in metastatic breast cancer cells as mentioned above. Moreover, we find differentially expressed tRNAs corresponding to codons previously reported to change KRAS protein levels when synonymously mutated (Lampson et al., 2013). Taken together, our results suggest that in order to increase KRAS translation efficiency it is not necessary to change the expression of multiple tRNAs but just that of a few specific ones. Particularly, we observe that codons corresponding to the tRNAs playing a role in these changes, are also codons enriched in the oncogenes BRAF, RAC1, RHOA and COL11A1. Our results suggest that certain tRNAs could be used as markers of oncogene-specific translation. Determination of tRNA abundance of different cell types may reveal previously unseen connections between translation and oncogene prevalence in cancer. It would also be interesting to investigate how tRNA modifications could also influence oncogene translation. Furthermore, Supek et al (Supek et al., 2014) show that selection acts on somatic synonymous mutations of oncogenes in tumor evolution. In many cases they are associated with changes in oncogene splicing in tumors. It would be interesting to further investigate if some of the recurrent synonymous mutations in those oncogenes correspond to changes towards enriching their coding sequence in proliferation-related codons, ultimately yielding to a greater translation efficiency.

The question remains as to the physiological role of this family codon bias. One possible explanation is that the protein levels of KRAS, BRAF, RAC1, RHOA and COL11A1 need to be tightly controlled. For example, during embryogenesis tRNA levels have been shown to vary in mouse brain and liver (Schmitt et al., 2014). Indeed, KRAS (Johnson et al., 1997), RAC1 (Sugihara et al., 1998; Corbetta et al., 2005; Roberts et al., 1999) and RHOA (Liu et al., 2001); (Hakem et al., 2005) are the only family members embryonically lethal in homozygous null mice. On the other hand, BRAF (Wojnowski et al., 1997) and COL11A1 (Li et al., 1995) homozygous mutants are also lethal together with RAF1 (Wojnowski et al., 1998) and COL5A1 (Wenstrup et al., 2004). Thus, cancer cells could take advantage of a developmental translation regulation to boost the translation of oncogenic transcripts to their own growth advantage.

Taken together, our work not only addresses a fundamental aspect of RAS biology but also provides insight into the controversial issue of how codon bias can influence protein expression. Collectively, our findings demonstrate that codon-driven translational efficiency can modulate protein expression of oncogenes in different cell contexts.

## Acknowledgments

We thank Eva Maria Novoa and Manuel Irimia for stimulating and critical discussions. We thank Disa Tehler for sending BJ/hTERT cells. We acknowledge the support of the Spanish Ministry of Economy, Industry and Competitiveness (MEIC) to the EMBL partnership, the Centro de Excelencia Severo Ochoa and the CERCA Programme / Generalitat de Catalunya.

## Author Contributions

Conceptualization, H.B., M.H.S. and L.S.; Methodology, H.B., M.H.S., M.W., L.S.; Software, M.H.S., M.W., X.H.A.; Experiments: H.B., Validation, H.B., M.H.S., M.W., Formal analysis, H.B., M.H.S., M.W, X.H.A.; Investigation, H.B., M.W., M.H.S.; Writing-Original Draft, H.B.; Writing-Review & Editing, H.B., M.H.S., M.W., X.H.A., L.S.; Visualization: H.B., M.W., M.H.S.; Funding Acquisition, L.S.; Supervision, M.H.S. and L.S.

## Methods

### Data Sources

#### Paralogs Ensembl

To define gene families we retrieved protein sequence similarity and family membership information from Ensembl. As we observed that Ensembl’s family classification often contained outliers with much lower sequence similarity compared to the other proteins, we applied another, more stringent filter: for each family we computed the similarity distribution of all members to a consensus member. We then removed all family members with a similarity of less than the mean similarity minus one standard deviation. We only considered families with at least three members.

### TCGA

Mutation data was obtained from The Cancer Genome Atlas (TCGA). We retrieved somatic mutations in coding regions for 20 cancer types: Bladder Urothelial Carcinoma, Breast invasive carcinoma, Cervical squamous cell carcinoma and endocervical adenocarcinoma, Colon adenocarcinoma, Glioblastoma multiforme, Head and Neck squamous cell carcinoma, Kidney renal clear cell carcinoma, Kidney renal papillary cell carcinoma, Acute Myeloid Leukemia, Brain Lower Grade Glioma, Liver hepatocellular carcinoma, Lung adenocarcinoma, Lung squamous cell carcinoma, Pancreatic adenocarcinoma, Pheochromocytoma and Paraganglioma, Prostate adenocarcinoma, Skin Cutaneous Melanoma, Stomach adenocarcinoma, Thyroid carcinoma, and Uterine Corpus Endometrial Carcinoma comprising a set of 5,960 samples.

### Cancer gene catalogue

We considered cancer driver genes to be those genes that had a significant (q < 0.01) number of non-silent mutations in at least 1 out of 21 cancer types in 4,742 patients as defined in Lawrence et al (Lawrence et al., 2014).

### Coding Sequences

The coding sequences of *Homo sapiens* were downloaded from the Consensus CDS (CCDS) project (ftp://ftp.ncbi.nlm.nih.gov/pub/CCDS/) release 2016/09/08. In the case of non-cancer genes, one unique canonical coding sequence was arbitrarily chosen for each protein based on Uniprot mapping to the CCDS. In the case of genes in the selected cancer gene families, the canonical coding sequence was chosen following the corresponding protein defined as canonical in Uniprot.

### GO gene sets

Gene ontology was downloaded as MySQL dump of the amiGO database release 2017/01, and human gene annotations were downloaded from amiGO database release 2018/01/04. We defined GO gene sets as follows: for each GO term, we retrieved all descendant GO terms (with any kind of relationship type) and assigned all associated genes. We selected all GO terms with a minimal distance to the root “biological process” term shorter or equal to 3, and at least 30 associated genes, resulting in a total of 708 gene sets. Note that there is a lot of overlap between these GO gene sets, with a protein appearing on average in 44 sets.

### Computational analysis

#### Codon usage PCA

We applied principal component analysis (PCA) to the relative synonymous codon frequencies (Sharp and Li, 1987) of all individual human coding sequences. Note that, contrary to other studies such as the one in Gingold et al (Gingold et al., 2014), we defined our PCA projection based on the codon usage distribution of individual genes, not of gene sets. By doing so, our projection is independent from the gene ontology annotations. In addition, PCA based on the average codon usage of gene sets may suffer from bias due to the fact that GO gene sets are highly overlapping. Thus, the codon usage of a specific gene may contribute to several gene set data points, which may in turn distort the real variation in codon usage. When computing the PCA of individual genes, we first excluded single codon families (AUG, UGG). In the case of coding sequences that lacked codons of a specific family (6.7% of total), we impute values with the average codon frequency across all genes. We applied the PCA projection to GO gene sets, by computing the mean of relative codon frequencies of all genes in the set.

#### Quantification of tRNA expression

tRNAseq mapping was performed using a specific pipeline for tRNAs (Hoffmann et al., 2018). The basic pipeline was adapted to paired-end sequencing data. Moreover, given that hydro-tRNA-seq yields short sequences, all reads over 10 nt were included after BBDuk adapter trimming. Isoacceptors were quantified as reads per million (RPM), summing up all reads mapping to isodecoders that share the same anticodon. Ambiguous reads mapping to genes of different isoacceptors were discarded. The human reference genome GRCh38 (GenBank 2339568) was used.

#### Relative codon usage

We correlated the relative codon usage of KRAS_WT_ and KRAS_HRAS_ (calculated by dividing each codon value by the sum of the codon values of a given amino acid). In order to be able to calculate the fold change we added a pseudo count to all values (+1). For the 4 other families we calculated this fold change of codon usage between the most mutated gene and the less mutated gene from the same family. We first performed a sequence alignment using TranslatorX (Abascal et al., 2010) to be able to compare only the codons that align between the two sequences. Finally, we calculated the relative codon usage and the fold change in the same way as done for the comparison between KRAS_WT_ and KRAS_HRAS_.

#### Differential tRNA anticodon abundance

We exclude anticodons for which there are no corresponding tRNA genes (Arg^GCG^, Gly^ACC^, His^ATG^, Leu^GAG^, Phe^AAA^, Thr^GGT^ and Val^GAC^) based on the tRNA gene prediction from the *H. sapiens* genome GRCh38/hg38 using tRNAscan-SE (Chan et al.). Next, we calculate the relative anticodon abundance: dividing each anticodon rpm value by the sum of the anticodons rpm values for a given amino acid). Differential relative expression analysis was performed using t-test, where p-values were FDR-corrected, with p < 0.05 as a cutoff.

### Sample preparation and experimental procedures

#### Cell lines

The cell lines included in this study were comprised of: HeLa, HEK293 and fibroblast BJ/hTERT (used in Gingold et al.(Gingold et al., 2014), kindly provided by the author Disa Tehler). Cells were maintained at 37 °C in a humidified atmosphere at 5% CO_2_ in DMEM 4.5g/L Glucose with UltraGlutamine media supplemented with 10% of Tet-free FBS (Clontech) and 1% penicillin/streptomycin.

#### Expression vector design

KRAS_HRAS_ was obtained from pBABE-Puro-KRas* (Addgene#46745). For conditional-gene overexpression experiments, KRAS_WT_ and KRAS_HRAS_ were cloned into a modified version of the XLone-GFP vector (Randolph et al., 2017) (Addgene#96930). The modification consisted of replacing the promoter of XLone-GFP with a bidirectional TRE3G promoter (Clontech) allowing the simultaneous expression of both KRAS genes. We use a FLAG tag and 3xHA to distinguish FLAG-KRAS_WT_ and 3xHA-KRAS_HRAS_ by size. We also inverted the tags FLAG-KRAS_HRAS_ and 3xHA-KRAS_WT._ The vector was co-transfected in different cell lines with the plasmid pCYL43 (Wang et al., 2008) containing the PiggyBac transposase. Cells were selected with blasticidin (HeLa: 5µg/mL, HEK293: 15µg/mL, BJ/hTERT: 5µg/mL). Gene expression was induced with doxycycline (HeLa: 100ng/mL, HEK293: 12ng/mL, BJ/hTERT: 500ng/mL).

#### Serum starvation assay

BJ/hTERT were grown in starvation media (1% Tet-free FBS) or non-starvation media (10% Tet-free FBS) for 48 hours. The expression of both KRAS_WT_ and KRAS_HRAS_ was measured after doxycycline induction overnight.

#### Cell lines assay

Established HeLa, HEK293 and BJ/hTERT cells were induced with doxycycline and the expression was measured after overnight incubation.

#### Cell growth

The cells were seeded at a density of 25000 cells per well in a 12-well plate and the counts were performed with Countess cell counting chamber slides and the Countess automated cell counter (ThermoFisher). The counts were carried out every 24 hours.

#### mRNA quantification

RNA isolation was performed with RNeasy kit (Qiagen). KRAS_WT_ and KRAS_HRAS_ transcript abundances were quantified by RT-qPCR (Power SYBR Green RNA-to-CT 1-Step Kit, ThermoFisher). Primers for FLAG-KRAS_WT_ amplification: forward 5’-CAAGGACGACGATGACAAG-3’ and reverse 5’-GAGAATATCCAAGAGACAGGTT-3’. Primers for 3xHA-KRAS_HRAS_ amplification: forward 5’-CCTGACTATGCGGGCTATC-3’ and reverse 5’-GGGTCGTATTCGTCCACAA-3’. As both genes are in the same expression cassette, for each sample, the Ct values for KRAS_WT_ were normalized to the KRAS_HRAS_, Δ*Ct*.= (*Ct*KRAS_WT_ - *Ct*KRAS_HRAS_) and represented as 2 ^- ΔCt^.

#### Quantitative protein blots

Cells were lysed using an M-PER buffer (ThermoFisher) supplemented with anti-proteases. Protein concentration was measured using a BCA Protein Assay Kit (Pierce). Equal amounts of each sample were mixed with 1x Laemmli buffer and boiled for 5 min. Samples were separated using 12% polyacrylamide gels (BioRad). Transfer was performed using the iBlot system (Invitrogen). Membranes were treated with Li-COR Odyssey blocking buffer for 1 hr at RT, then incubated with primary antibody (1:1000) in 0.2% Tween-20/Li-COR odyssey blocking buffer overnight at 4°C. Following three 5 min washes in TBS-T, the membrane was incubated with secondary antibodies (1:10000) in 0.2% Tween-20/Li-COR Odyssey blocking buffer for 45 min at RT. Following three 5 min washes in TBS-T, the membrane was scanned using the Li-COR Odyssey Imaging System. We used the following primary antibodies: anti-pan-RAS (Abcam, ab52939) and anti-*β*-actin (Sigma, A2228) and were detected using a goat anti-rabbit (Abcam, ab216773) or goat anti-mouse (Abcam, ab216776) IgG antibody conjugated to an IRdye at 800CW and 680CW, respectively. Visualization and quantification was done using ImageJ and Image Studio Lite (LI-COR).

#### Hydro-tRNA sequencing

Total RNA from HEK293 and HeLa was extracted using the miRNeasy Mini kit. For each sample, 20 µg of total RNA was treated following the protocol of hydro-tRNAseq (Gogakos et al., 2017). Sequencing was done on Illumina HiSeq 2500 platform in 50bp paired-end format. Raw data have been deposited in the ArrayExpress database (Kolesnikov et al., 2015) at EMBL-EBI (www.ebi.ac.uk/arrayexpress) under accession number E-MTAB-8144.

## Supplemental figure legends

**Figure S1:**
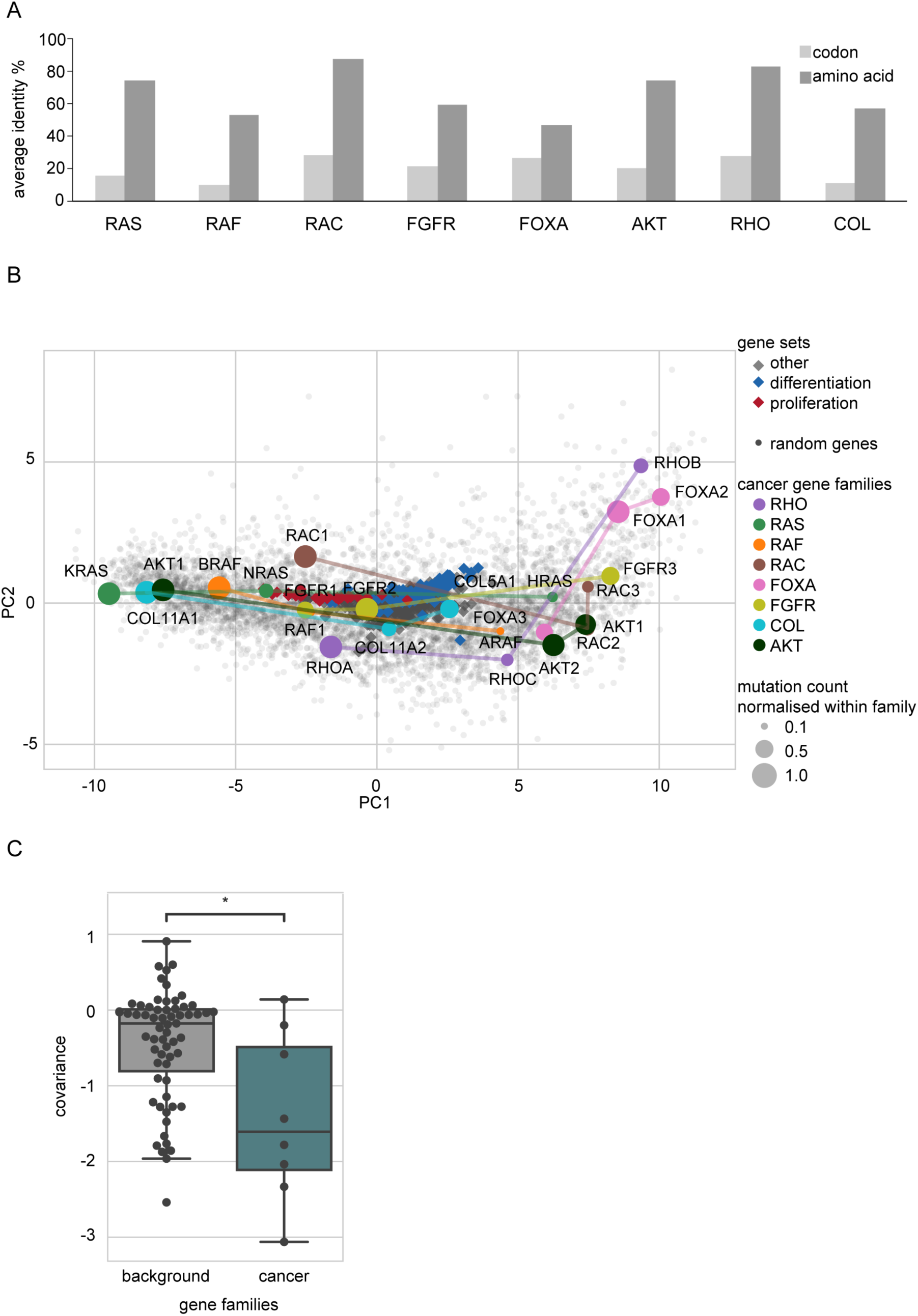
Association between codon usage and mutation frequency in genes from 8 different families, related to Figure 1. A. Codon and amino acid identity of gene families. B. PCA projection of the human codon usage. The location of each gene is determined by its codon usage. C. Distribution of the covariance of mutation count normalized within family and PC1. Covariance is significantly more negative for cancer gene families than for the background non-cancer related gene families (W.M.W. test p<0.018). In particular, the covariance is negative for 7 (RAS, RAF, RHO, RAC, FGFR, AKT and COL) out of 8 cancer gene families.

**Figure S2:**
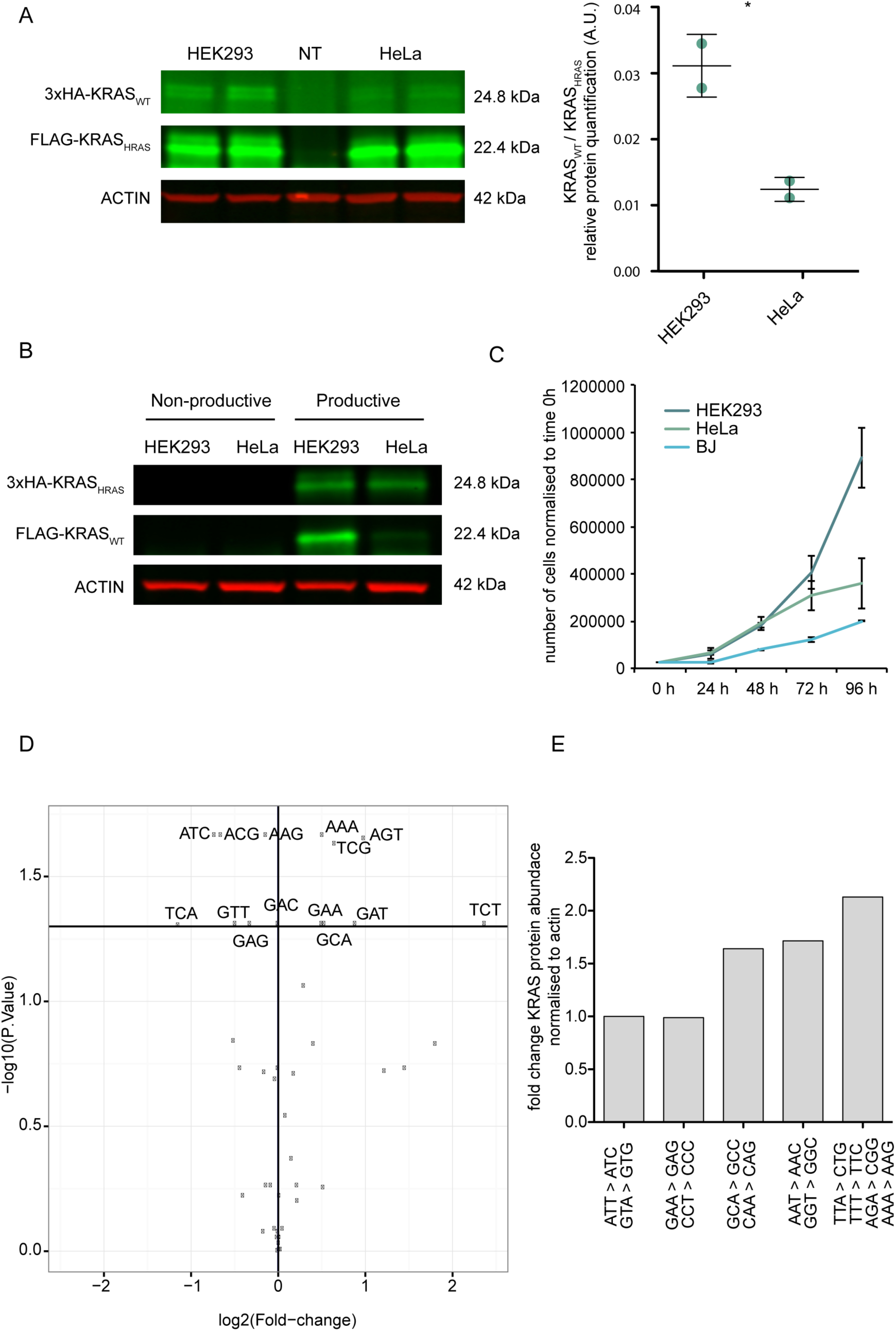
Control with inverted tags, cell lines growth curves and differential tRNAs expression, related to Figure 3. A. Western blot analysis of the levels of KRAS_WT_ and KRAS_HRAS_ with inverted tags in HEK293 and HeLa cells. The protein ratio KRAS_WT_/KRAS_HRAS_ varies between the different cell lines similarly, regardless of the tag. Error bars in A represent SEM of two independent experiments *, p < 0.05. B. Comparison of expression between the productive and non-productive (without RBS and ATG) expression cassette. No KRAS_WT_ or KRAS_HRAS_ expression is observed when translation is suppressed. C. HEK293, HeLa and BJ/hTERT cell count over 96 hours. D. Volcano plot showing relative tRNA differential expression in log2 fold change between HEK293 and HeLa. E. Immunoblot data taken and quantified from Lampson et al (Lampson et al., 2013) where KRAS codons were progressively converted.

**Figure S3:**
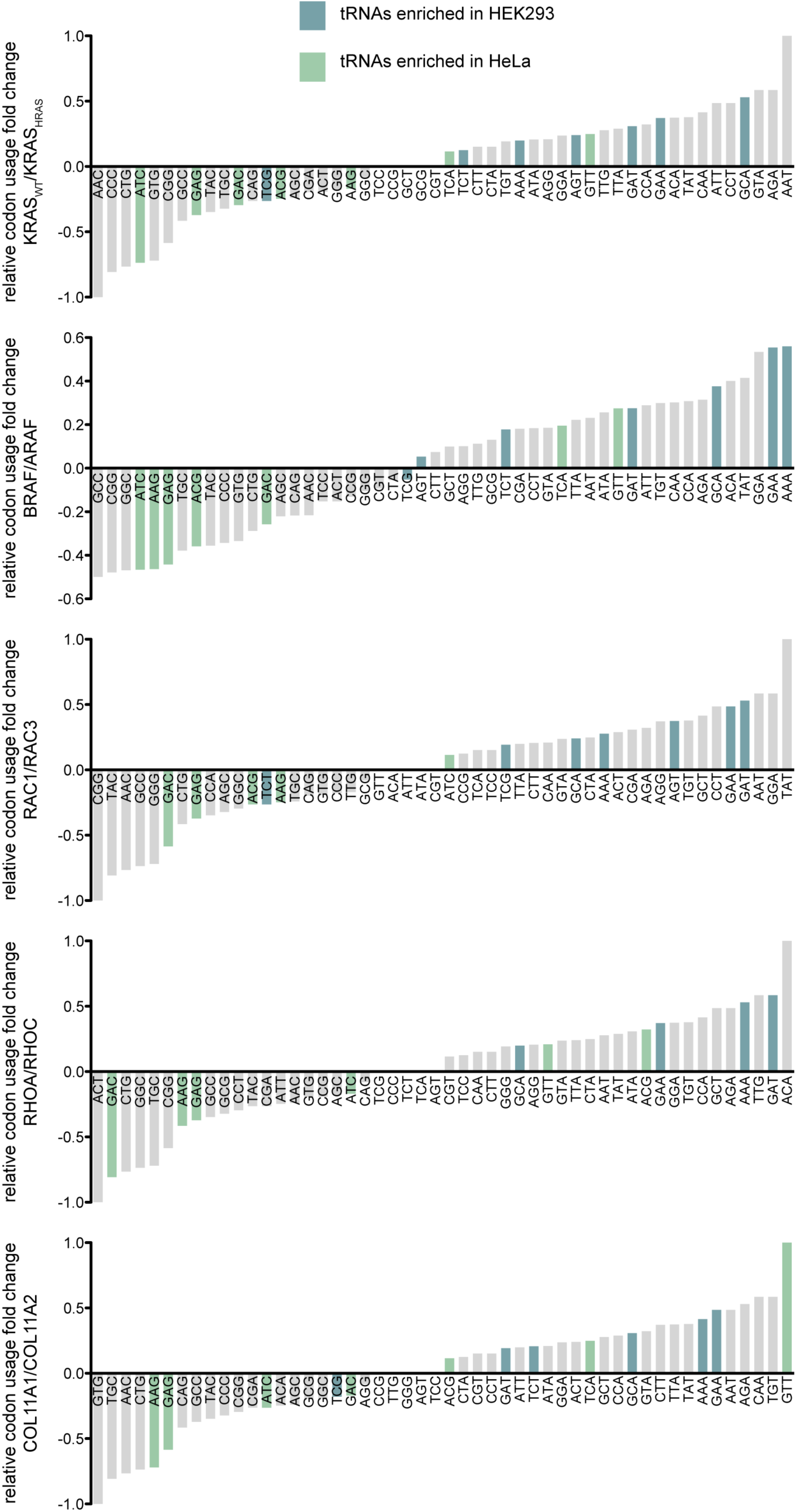
Relative codon usage between the most frequently and the less frequently mutated gene, related to Figure 3 and 4. Fold change of the relative codon abundance (pseudocount +1) between the most frequently and the less frequently mutated gene for the 4 families displaying the highest negative covariance together with the RAS family. tRNAs differentially expressed between HEK293 (blue) and HeLa (green). Generally, the tRNAs enriched in HEK293 and matching the codons enriched in KRAS_WT_ are enriched in BRAF, RAC1, RHOA and COL11A1 and vice versa with the tRNAs enriched in HeLa.

**Figure S4:**
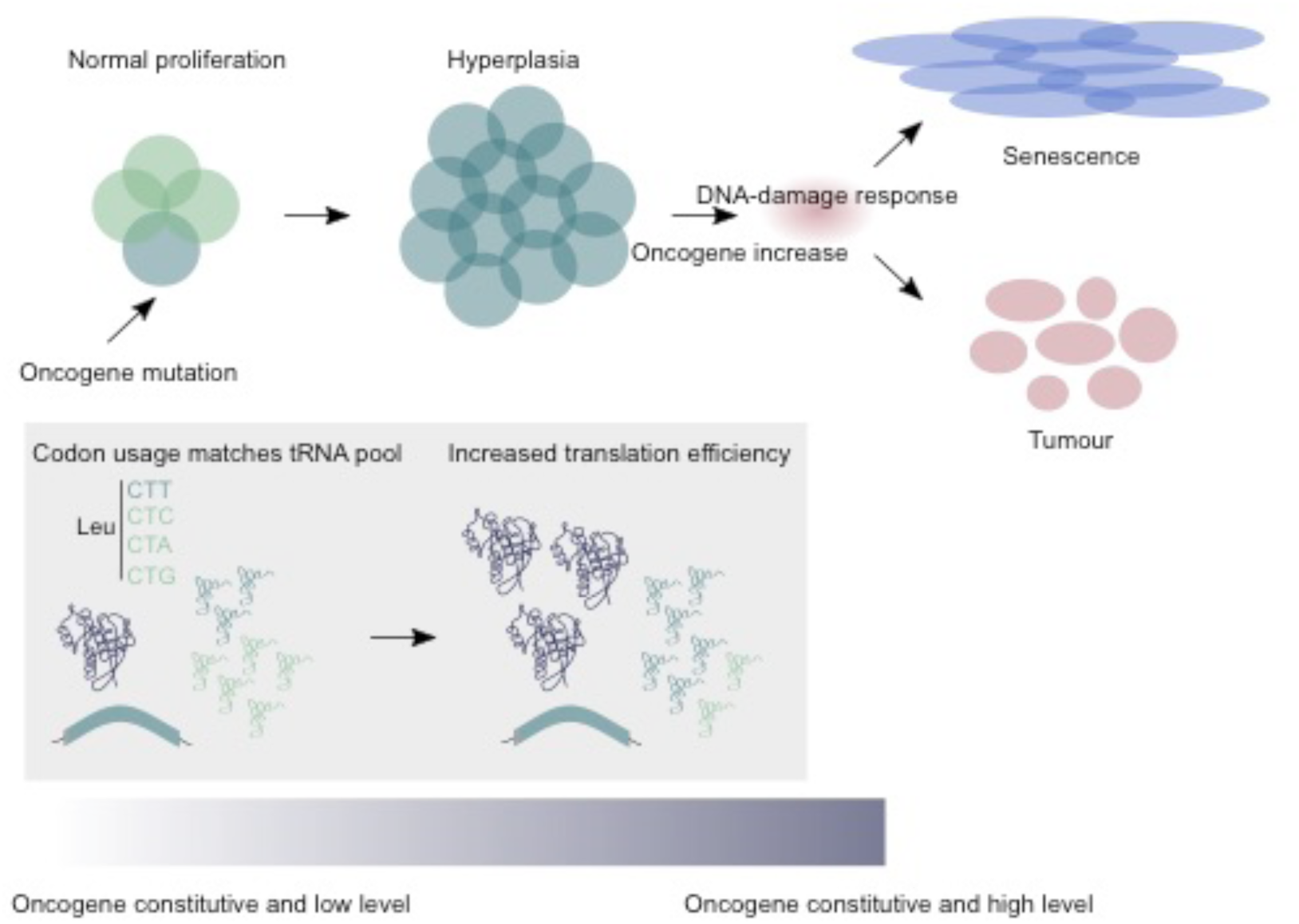
Based on the three-stage carcinogenesis model for RAS-induced tumours proposed in Sarkisian et al (Sarkisian et al., 2007). Adapted figure from “Barriers to RAS transformation”, Ferbeyre G, Nat Cell Biol, 2007 (Ferbeyre, 2007). Oncogenic mutations lead to a constitutive activation of the oncogene that promotes cell proliferation. Low levels of oncogene are not transforming. An increase of oncogene translation efficiency will increase oncogene levels. High oncogene levels and activity would transform cells if cells evade senescence.

**Table S1: Data concerning the genes from the 8 different families, related to Figure 1 and S1**.

**Table S2: Data concerning background and cancer gene families, related to Figure 1 and S1**.

**Table S3: tRNA expression (rpm) data processed, related to Figure 3**.

**Table S4: tRNA expression relative fold change between HEK293 and HeLa and relative codon abundance for KRAS**_**WT**_ **and KRAS**_**HRAS**_, **related to Figure 3 and Figure S2D**.

## References

Abascal, F., Zardoya, R., and Telford, M.J. (2010). TranslatorX: multiple alignment of nucleotide sequences guided by amino acid translations. Nucleic Acids Res. 38, W7–W13.

Apolloni, A., Prior, I.A., Lindsay, M., Parton, R.G., and Hancock, J.F. (2000). H-ras but Not K-ras Traffics to the Plasma Membrane through the Exocytic Pathway. Mol. Cell. Biol. 20, 2475–2487.

Barbieri, C.E., Baca, S.C., Lawrence, M.S., Demichelis, F., Blattner, M., Theurillat, J.-P., White, T.A., Stojanov, P., Van Allen, E., Stransky, N., et al. (2012). Exome sequencing identifies recurrent SPOP, FOXA1 and MED12 mutations in prostate cancer. Nat. Genet. 44, 685–689.

Bodemann, B.O., and White, M.A. (2013). Ras GTPases: Codon Bias Holds KRas Down but Not Out. Curr. Biol. 23, R17–R20.

Boël, G., Letso, R., Neely, H., Nicholson Price, W., Wong, K.-H., Su, M., Luff, J.D., Valecha, M., Everett, J.K., Acton, T.B., et al. (2016). Codon influence on protein expression in E. coli correlates with mRNA levels. Nature 529, 358–363.

Chan, P.P., Lin, B.Y., Mak, A.J., and Lowe, T.M. tRNAscan-SE 2.0: Improved Detection and Functional Classification of Transfer RNA Genes.

Corbetta, S., Gualdoni, S., Albertinazzi, C., Paris, S., Croci, L., Consalez, G.G., and de Curtis, I. (2005). Generation and characterization of Rac3 knockout mice. Mol. Cell. Biol. 25, 5763–5776.

Dittmar, K.A., Goodenbour, J.M., and Pan, T. (2006). Tissue-specific differences in human transfer RNA expression. PLoS Genet. 2, e221.

Drosten, M., Simón-Carrasco, L., Hernández-Porras, I., Lechuga, C.G., Blasco, M.T., Jacob, H.K.C., Fabbiano, S., Potenza, N., Bustelo, X.R., Guerra, C., et al. (2017). H-Ras and K-Ras Oncoproteins Induce Different Tumor Spectra When Driven by the Same Regulatory Sequences. Cancer Res. 77, 707–718.

Ferbeyre, G. (2007). Barriers to Ras transformation. Nat. Cell Biol. 9, 483–485.

Fu, J., Dang, Y., Counter, C., and Liu, Y. (2018). Codon usage regulates human KRAS expression at both transcriptional and translational levels. J. Biol. Chem. 293, 17929–17940.

Gingold, H., Tehler, D., Christoffersen, N.R., Nielsen, M.M., Asmar, F., Kooistra, S.M., Christophersen, N.S., Christensen, L.L., Borre, M., Sørensen, K.D., et al. (2014). A dual program for translation regulation in cellular proliferation and differentiation. Cell 158, 1281–1292.

Gogakos, T., Brown, M., Garzia, A., Meyer, C., Hafner, M., and Tuschl, T. (2017). Characterizing Expression and Processing of Precursor and Mature Human tRNAs by Hydro-tRNAseq and PAR-CLIP. Cell Rep. 20, 1463–1475.

Goodarzi, H., Nguyen, H.C.B., Zhang, S., Dill, B.D., Molina, H., and Tavazoie, S.F. (2016). Modulated Expression of Specific tRNAs Drives Gene Expression and Cancer Progression. Cell 165, 1416–1427.

Haigis, K.M., Kendall, K.R., Wang, Y., Cheung, A., Haigis, M.C., Glickman, J.N., Niwa-Kawakita, M., Sweet-Cordero, A., Sebolt-Leopold, J., Shannon, K.M., et al. (2008). Differential effects of oncogenic K-Ras and N-Ras on proliferation, differentiation and tumor progression in the colon. Nat. Genet. 40, 600–608.

Hakem, A., Sanchez-Sweatman, O., You-Ten, A., Duncan, G., Wakeham, A., Khokha, R., and Mak, T.W. (2005). RhoC is dispensable for embryogenesis and tumor initiation but essential for metastasis. Genes Dev. 19, 1974–1979.

Hanson, G., and Coller, J. (2018). Codon optimality, bias and usage in translation and mRNA decay. Nat. Rev. Mol. Cell Biol. 19, 20–30.

Hobbs, G.A., Aaron Hobbs, G., Der, C.J., and Rossman, K.L. (2016). RAS isoforms and mutations in cancer at a glance. J. Cell Sci. 129, 1287–1292.

Hoffmann, A., Fallmann, J., Vilardo, E., Mörl, M., Stadler, P.F., and Amman, F. (2018). Accurate mapping of tRNA reads. Bioinformatics 34, 2339.

Johnson, L., Greenbaum, D., Cichowski, K., Mercer, K., Murphy, E., Schmitt, E., Bronson, R.T., Umanoff, H., Edelmann, W., Kucherlapati, R., et al. (1997). K-ras is an essential gene in the mouse with partial functional overlap with N-ras. Genes Dev. 11, 2468–2481.

Koera, K., Nakamura, K., Nakao, K., Miyoshi, J., Toyoshima, K., Hatta, T., Otani, H., Aiba, A., and Katsuki, M. (1997). K-ras is essential for the development of the mouse embryo. Oncogene 15, 1151–1159.

Kolesnikov, N., Hastings, E., Keays, M., Melnichuk, O., Tang, Y.A., Williams, E., Dylag, M., Kurbatova, N., Brandizi, M., Burdett, T., et al. (2015). ArrayExpress update--simplifying data submissions. Nucleic Acids Res. 43, D1113–D1116.

Lampson, B.L., Pershing, N.L.K., Prinz, J.A., Lacsina, J.R., Marzluff, W.F., Nicchitta, C.V., MacAlpine, D.M., and Counter, C.M. (2013). Rare codons regulate KRas oncogenesis. Curr. Biol. 23, 70–75.

Lawrence, M.S., Stojanov, P., Mermel, C.H., Robinson, J.T., Garraway, L.A., Golub, T.R., Meyerson, M., Gabriel, S.B., Lander, E.S., and Getz, G. (2014). Discovery and saturation analysis of cancer genes across 21 tumour types. Nature 505, 495–501.

Li, Y., Lacerda, D.A., Warman, M.L., Beier, D.R., Yoshioka, H., Ninomiya, Y., Oxford, J.T., Morris, N.P., Andrikopoulos, K., and Ramirez, F. (1995). A fibrillar collagen gene, Col11a1, is essential for skeletal morphogenesis. Cell 80, 423–430.

Liu, A.X., Rane, N., Liu, J.P., and Prendergast, G.C. (2001). RhoB is dispensable for mouse development, but it modifies susceptibility to tumor formation as well as cell adhesion and growth factor signaling in transformed cells. Mol. Cell. Biol. 21, 6906–6912.

Newman, Z.R., Young, J.M., Ingolia, N.T., and Barton, G.M. (2016). Differences in codon bias and GC content contribute to the balanced expression of TLR7 and TLR9. Proc. Natl. Acad. Sci. U. S. A. 113, E1362–E1371.

Nowell, P. (1976). The clonal evolution of tumor cell populations. Science 194, 23–28.

Omerovic, J., Hammond, D.E., Clague, M.J., and Prior, I.A. (2008). Ras isoform abundance and signalling in human cancer cell lines. Oncogene 27, 2754–2762.

Pavon-Eternod, M., Gomes, S., Geslain, R., Dai, Q., Rosner, M.R., and Pan, T. (2009). tRNA over-expression in breast cancer and functional consequences. Nucleic Acids Res. 37, 7268–7280.

Pershing, N.L.K., Lampson, B.L., Belsky, J.A., Kaltenbrun, E., MacAlpine, D.M., and Counter, C.M. (2015). Rare codons capacitate Kras-driven de novo tumorigenesis. J. Clin. Invest. 125, 222–233.

Potenza, N., Vecchione, C., Notte, A., De Rienzo, A., Rosica, A., Bauer, L., Affuso, A., De Felice, M., Russo, T., Poulet, R., et al. (2005). Replacement of K-Ras with H-Ras supports normal embryonic development despite inducing cardiovascular pathology in adult mice. EMBO Rep. 6, 432–437.

Presnyak, V., Alhusaini, N., Chen, Y.-H., Martin, S., Morris, N., Kline, N., Olson, S., Weinberg, D., Baker, K.E., Graveley, B.R., et al. (2015). Codon Optimality Is a Major Determinant of mRNA Stability. Cell 160, 1111–1124.

Prior, I.A., Muncke, C., Parton, R.G., and Hancock, J.F. (2003). Direct visualization of Ras proteins in spatially distinct cell surface microdomains. J. Cell Biol. 160, 165–170.

Quinlan, M.P., and Settleman, J. (2008). Explaining the preponderance of Kras mutations in human cancer: An isoform-specific function in stem cell expansion. Cell Cycle 7, 1332–1335.

Radhakrishnan, A., Chen, Y.-H., Martin, S., Alhusaini, N., Green, R., and Coller, J. (2016). The DEAD-Box Protein Dhh1p Couples mRNA Decay and Translation by Monitoring Codon Optimality. Cell 167, 122–132.e9.

Randolph, L.N., Bao, X., Zhou, C., and Lian, X. (2017). An all-in-one, Tet-On 3G inducible PiggyBac system for human pluripotent stem cells and derivatives. Sci. Rep. 7, 1549.

Roberts, A.W., Kim, C., Zhen, L., Lowe, J.B., Kapur, R., Petryniak, B., Spaetti, A., Pollock, J.D., Borneo, J.B., Bradford, G.B., et al. (1999). Deficiency of the hematopoietic cell-specific Rho family GTPase Rac2 is characterized by abnormalities in neutrophil function and host defense. Immunity 10, 183–196.

Rocks, O., Peyker, A., and Bastiaens, P.I.H. (2006). Spatio-temporal segregation of Ras signals: one ship, three anchors, many harbors. Curr. Opin. Cell Biol. 18, 351–357.

Sarkisian, C.J., Keister, B.A., Stairs, D.B., Boxer, R.B., Moody, S.E., and Chodosh, L.A. (2007). Dose-dependent oncogene-induced senescence in vivo and its evasion during mammary tumorigenesis. Nat. Cell Biol. 9, 493–505.

Schmitt, B.M., Rudolph, K.L.M., Karagianni, P., Fonseca, N.A., White, R.J., Talianidis, I., Odom, D.T., Marioni, J.C., and Kutter, C. (2014). High-resolution mapping of transcriptional dynamics across tissue development reveals a stable mRNA-tRNA interface. Genome Res. 24, 1797–1807.

Schroeder, M.P., Rubio-Perez, C., Tamborero, D., Gonzalez-Perez, A., and Lopez-Bigas, N. (2014). OncodriveROLE classifies cancer driver genes in loss of function and activating mode of action. Bioinformatics 30, i549–i555.

Sharp, P.M., and Li, W.H. (1987). The codon Adaptation Index--a measure of directional synonymous codon usage bias, and its potential applications. Nucleic Acids Res. 15, 1281–1295.

Sobhani, N., Ianza, A., D’Angelo, A., Roviello, G., Giudici, F., Bortul, M., Zanconati, F., Bottin, C., and Generali, D. (2018). Current Status of Fibroblast Growth Factor Receptor-Targeted Therapies in Breast Cancer. Cells 7.

Sugihara, K., Nakatsuji, N., Nakamura, K., Nakao, K., Hashimoto, R., Otani, H., Sakagami, H., Kondo, H., Nozawa, S., Aiba, A., et al. (1998). Rac1 is required for the formation of three germ layers during gastrulation. Oncogene 17, 3427–3433.

Supek, F. (2016). The Code of Silence: Widespread Associations Between Synonymous Codon Biases and Gene Function. J. Mol. Evol. 82, 65–73.

Supek, F., Miñana, B., Valcárcel, J., Gabaldón, T., and Lehner, B. (2014). Synonymous mutations frequently act as driver mutations in human cancers. Cell 156, 1324–1335.

Torrent, M., Chalancon, G., de Groot, N.S., Wuster, A., and Madan Babu, M. (2018). Cells alter their tRNA abundance to selectively regulate protein synthesis during stress conditions. Sci. Signal. 11.

Wang, W., Lin, C., Lu, D., Ning, Z., Cox, T., Melvin, D., Wang, X., Bradley, A., and Liu, P. (2008). Chromosomal transposition of PiggyBac in mouse embryonic stem cells. Proc. Natl. Acad. Sci. U. S. A. 105, 9290–9295.

Wenstrup, R.J., Florer, J.B., Brunskill, E.W., Bell, S.M., Chervoneva, I., and Birk, D.E. (2004). Type V collagen controls the initiation of collagen fibril assembly. J. Biol. Chem. 279, 53331–53337.

Wojnowski, L., Zimmer, A.M., Beck, T.W., Hahn, H., Bernal, R., Rapp, U.R., and Zimmer, A. (1997). Endothelial apoptosis in Braf-deficient mice. Nat. Genet. 16, 293–297.

Wojnowski, L., Stancato, L.F., Zimmer, A.M., Hahn, H., Beck, T.W., Larner, A.C., Rapp, U.R., and Zimmer, A. (1998). Craf-1 protein kinase is essential for mouse development. Mech. Dev. 76, 141–149.

Wu, Q., Medina, S.G., Kushawah, G., DeVore, M.L., Castellano, L.A., Hand, J.M., Wright, M., and Bazzini, A.A. (2019). Translation affects mRNA stability in a codon-dependent manner in human cells. eLife 8.

Yan, J., Roy, S., Apolloni, A., Lane, A., and Hancock, J.F. (1998). Ras isoforms vary in their ability to activate Raf-1 and phosphoinositide 3-kinase. J. Biol. Chem. 273, 24052–24056.

